# Plasma miRNA-214 is a predictive candidate biomarker of progression speed in patients with ALS

**DOI:** 10.1101/2022.05.04.490596

**Authors:** Min-Young Noh, Min-Soo Kwon, Ki-Wook Oh, Minyeop Nahm, Jinseok Park, Young-Eun Kim, Hee Kyung Jin, Jae-sung Bae, Chang-Seok Ki, Seung Hyun Kim

**Author notes:** **Corresponding author:** Prof. Seung H Kim, M.D., Ph.D., Corresponding author’s address: Director of Cell Therapy Center, Department of Neurology, College of Medicine, Hanyang University, 222-1, Wangsimni-ro, Seongdong-gu, Seoul, 04763, Republic of Korea, Corresponding author’s; Corresponding author’s. These authors contributed equally to the manuscript (Min-Young Noh, Min-Soo Kwon).

## Abstract

This study was designed to develop and validate a reliable biomarker to predict the progression speed reflecting immune function of amyotrophic lateral sclerosis (ALS). After establishing the induced microglia model (iMGs) derived from peripheral blood monocytes, comparative studies to find factors related to phagocytic differences between iMGs of patients with rapidly progressive ALS [ALS(R)-iMGs, n = 15] and those of patients with slowly progressive ALS [ALS(S)-iMGs, n = 14] were conducted in the discovery cohort. To validate discovered candidate and whether it could be used as a reliable biomarker predicting the progression speed of ALS, we recruited 132 patients with ALS and 30 age-matched healthy controls as the validation cohort. ALS(R)-iMGs showed impaired phagocytic function. Transcriptomic analysis revealed that the perturbed phagocytosis in ALS(R)-iMGs was related to the decreased expression of NCKAP1 (NCK-associated protein 1) and NCKAP1 overexpression rescued the impaired phagocytic function. miRNA-214-3p targeting NCKAP1 in ALS-iMGs was correlated with progression speed in the discovery cohort. The validation cohort revealed that plasma miRNA-214-3p levels were significantly increased in ALS patients (*p* < 0.0001, AUC = 0.839), correlated with disease progression speed (*p* = 0.0005), and distinguished the rapidly progressive subgroup (Q1) from the slowly progressive (Q4, *p* = 0.029), respectively. Plasma miRNA-214-3p can predict the progression speed in ALS. Plasma miRNA-214-3p could be used as a simple and easily accessible biomarker for predicting the future progression speed related to phagocytic dysfunction in ALS patients.

## INTRODUCTION

Amyotrophic lateral sclerosis (ALS) is a fatal neurodegenerative disease characterized by the loss of motor neurons and inflammation in the motor neural axis, including the primary motor cortex, brainstem, and spinal cord, resulting in muscle weakness, wasting, respiratory paralysis, and, ultimately, death within 3 – 5 years (1).

The recent revolution in molecular biology has led to an exponential increase in attempts to develop prognostic and therapeutic decisional markers such as neurofilament light chain (NfL), TAR DNA-binding protein 43 (TDP-43), and neurotrophin receptor p75 (2-4). However, considering the genetic and clinical heterogeneity of ALS, there are still limited biomarkers that can be applied to clinical practice. Lessons from previous clinical trials in ALS have led us to develop reliable biomarkers that stratify patients’ groups (5). With the concept of a companion diagnostic and a complementary diagnostic in clinical trials (6, 7), the discovery of robust and well-validated new biomarkers would be helpful to the therapeutic decision-making process, enrolling the appropriate participants and monitoring the response of treatment. The US Food and Drug Administrations (FDA) acknowledged that the judicious use of biomarkers is expected to play an essential role in minimizing the risk of clinical trial failure by enriching trial populations with specific molecular subtypes responding better to the tested therapies (8). There is an ongoing unmet need for biomarkers that can reliably and efficiently stratify patients according to anticipated progression status, distinguish between responders and non-responders based on the mode of action (9), and help optimize treatment decisions in clinical trials.

Reactive microglial cells play a crucial role in determining the clinical progression of ALS (10). However, a previous clinical trial suggested that complete suppression of microglial function seems to be irrelevant to optimal control of neuro-inflammatory processes (11, 12). Thus, therapeutic strategies targeting microglia or immune functions should be tailored to their affected functional pathways according to progression speed rather than global suppression. Our *a priori* assumption was that the progression speed of ALS might be associated with the microglia phagocytic function, which was indirectly suggested by animal model and human data (13-15). We hypothesized that microglial phagocytic functions differ according to the speed of progression of ALS. This study aimed to discover and validate a reliable biomarker to predict progression speed reflecting immune function in ALS patients.

After establishing the induced microglia (iMGs) model using peripheral blood monocytes (PBMC) described in previous studies (16-18), serial comparative studies delineating the molecular and functional differences in iMGs, including gene expression and phagocytic function, were conducted in slowly and rapidly progressive patients with ALS. Phagocytic dysfunction was remarkable in iMGs of ALS(R). By transcriptomic analysis and web-based prediction algorithms, we found candidate biomarkers involved in the phagocytic dysfunction by rapid progression. After internal validation of the candidate biomarkers in 29 ALS patients (discovery cohort), we validated its significance in a large validation cohort (132 ALS patients and 30 age-matched healthy control (HC)) and whether it could be used as a reliable biomarker predicting progression speed in ALS.

## RESULTS

### ALS(R)-iMGs show impaired phagocytic function

In the discovery study, the mean ΔFS (T_0_ -T_2_) was 1.6 in ALS(R) and 0.2 in ALS(S) (Table 1 and Supplemental Table 1). The mean ΔFS (T_0_-T_2_) disease duration was 13.7 months in ALS(R) and 53.0 months in ALS(S). Age at symptom onset (T_0_) was younger in ALS(S), but the age at the iMG sampling time (T_2_) was similar between the two groups. The mean difference between ALSFRS-R and ΔFS (T_0_-T_2_) in the two groups was 10.2 (*p* < 0.001), 1.4 points/months (*p* < 0.001), suggesting that clinical status at T_2_ had deteriorated more in the ALS(R) group. ALSFRS-R measured at T_2_ was 27.7 in ALS(R) and 37.9 in ALS(S). Plasma cytokines were also different between the two groups. Plasma IL-8 level was increased in ALS(R), consistent with a previous study (19) (Table 1, Supplemental Table 5).

**Table 1.**
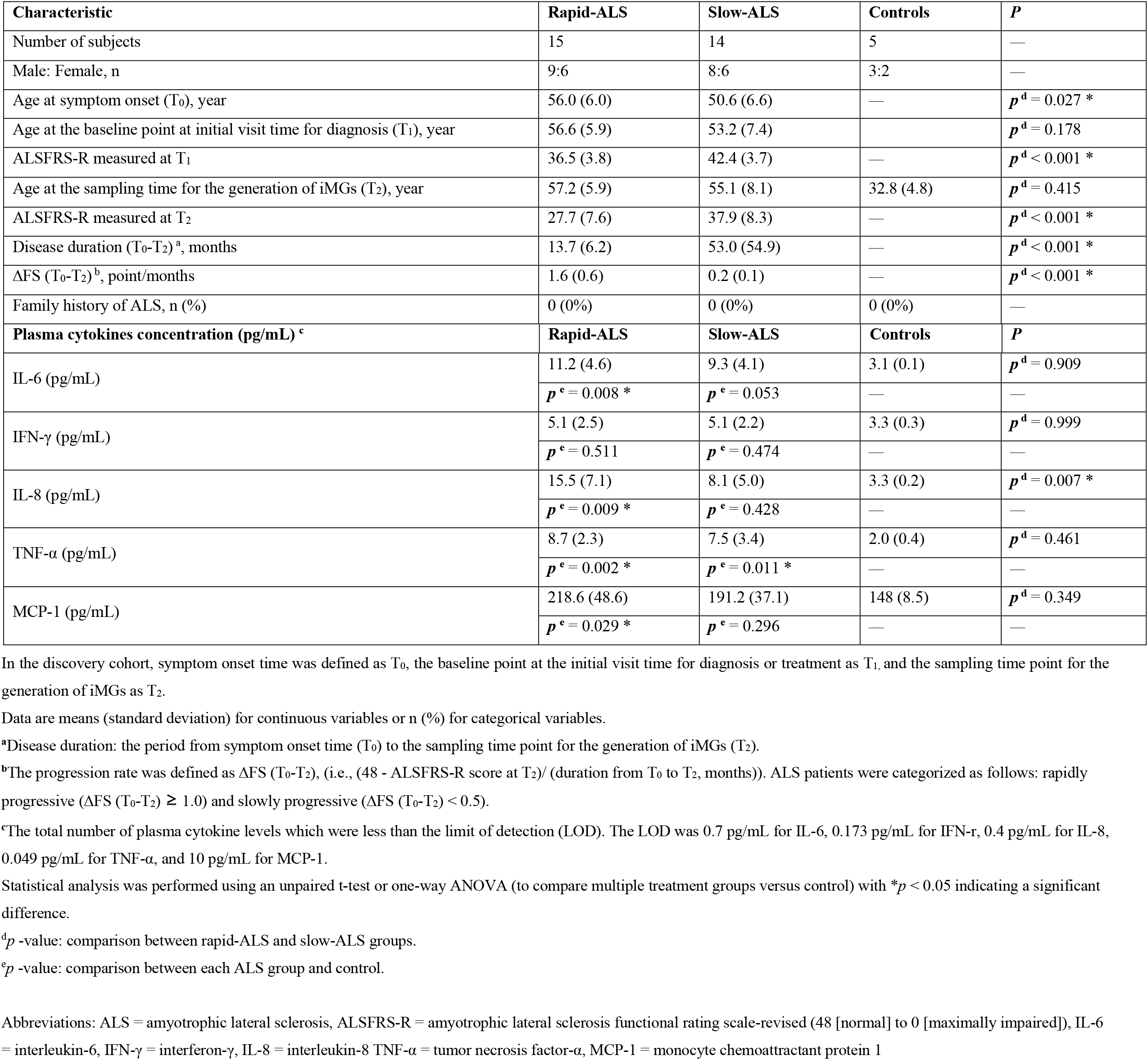
Demographics and characteristics of the discovery cohort.

The monocytes of each HC were treated with GM-CSF (10 ng/mL) and IL-34 (100 ng/mL) for 21 days to induce microglia-like cell branched morphology (iMGs). The cytokine cocktail shifted the cell population toward CD11b^+^CD45^+^ (Figure 1A) and expressed highly resident microglial signature genes, *P2RY12, OLFML3, TGFBR1, TREM2*, and *TMEM119* (Figure 1B) (20), consistent with previous studies that iMGs could recapitulate the main signatures of microglia (16).

**Figure 1:**
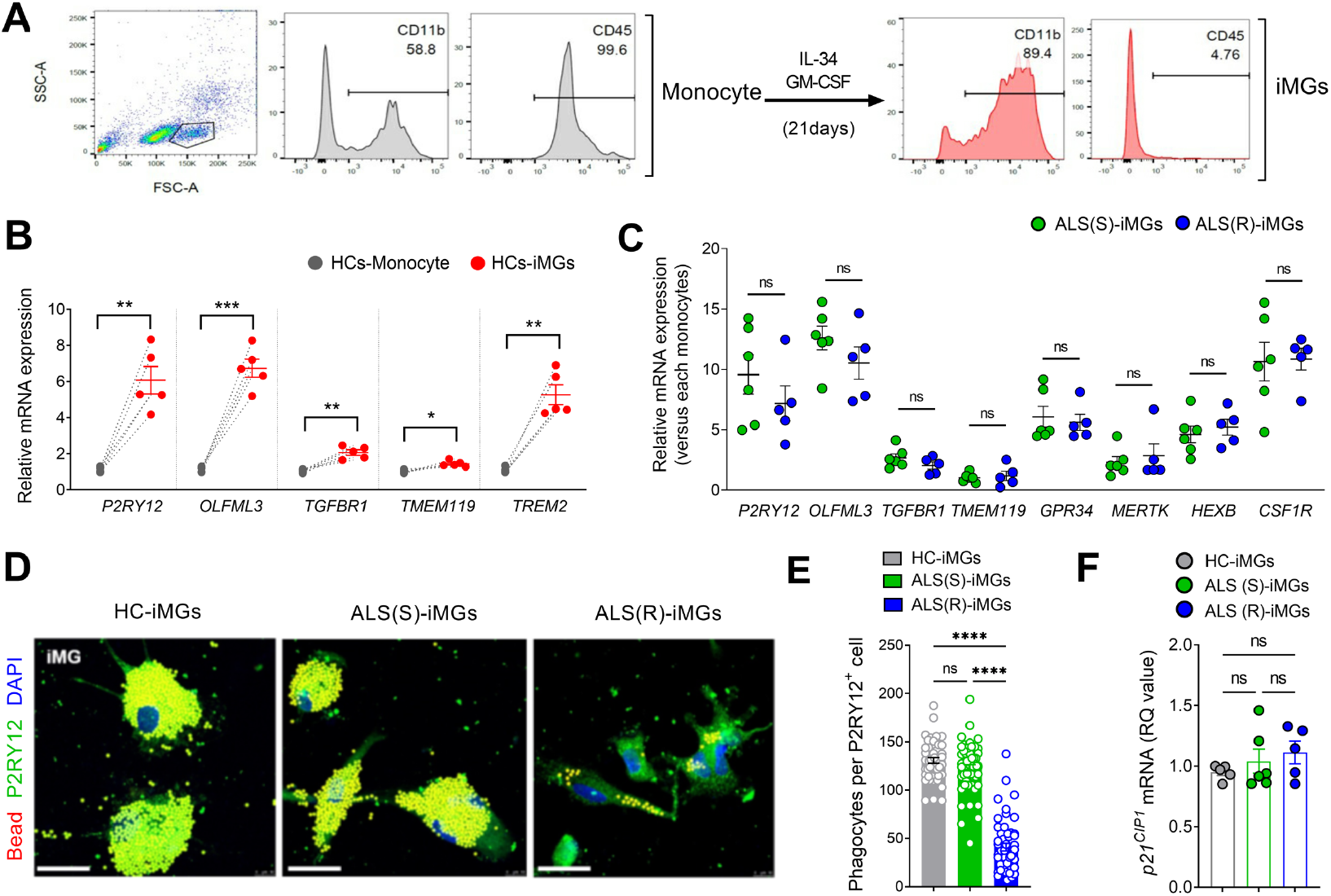
Impaired phagocytic function of ALS(R)-iMGs. **(A)** Representative flow cytometry plots showing microglia (CD45^lo^ CD11b^+^) from iMGs generated by treating monocytes with GM-CSF and IL-34 for 21 days. **(B)** Microglial signature gene expression (*P2RY12, OLFML3, TGFBR1, TMEM119*, and *TREM2*) of iMGs analyzed by RT-qPCR. Each dot represents data from individual-subject-derived HC-iMGs and monocytes (n = 5). **(C)** The expression of major homeostatic microglial signature genes from ALS(R)-iMGs (n = 5) and ALS(S)-iMGs (n = 6) by RT-qPCR. Each dot represents data from individual-subject-derived iMGs. **(D)** Representative confocal images of phagocytic activity from iMGs from each group incubated with latex beads for 24 h and immunostained with P2RY12, as iMG markers (green). **(E)** Mean number of phagocytosed beads per P2RY12^+^ cell per field quantified as in (**D**). Quantification of at least 50 cells from 10 iMGs per subject was performed (HC-iMGs: n = 5; ALS(R)-iMGs: n = 5; ALS(S)-iMGs: n = 6). **(F)** RT-qPCR analysis of cell senescence marker, *p21* expression in the three groups of iMGs. Each data point represents individual-subject-derived iMGs. Data were first tested for normality test. Statistical analysis between two groups was performed using Paired t-test (B) or Unpaired t-test (C) and between multiple groups using Kruskal-Wallis with post-hoc Dunn’s analysis (E) or one-way ANOVA with Tukey analysis (F). Values are the mean ± SEM (*P < 0.05, **P < 0.01, ***P < 0.001, ****P < 0.0001, ns: not significant). Scale bar: 25 μm.

For serial comparative studies, we generated iMGs from ALS(R) (R1 – R5) and ALS(S) (S1 – S6) patients. Both ALS(S)-iMGs and ALS(R)-iMGs significantly expressed microglia signature genes (*P2RY12, OLFML3, TGFBR1, GPR34, MERTK, HEXB, CSF1R, and TMEM119*) (20), which were rarely expressed by monocytes (Figure 1C). The most significant finding was that phagocytic function was severely impaired in ALS(R)-iMGs, whereas ALS(S)-iMGs exhibited no significant differences in phagocytic function compared to HC-iMGs (Figure 1D, E). However, defective phagocytic function was not associated with cellular senescence, with no difference in the expression of cellular senescence markers *p21*^*CIP1*^ (Figure 1F). Thus, we ruled out aging as a factor in the phagocytic dysfunction of ALS(R)-iMGs.

### Defective phagocytosis in ALS(R)-iMGs is associated with decreased NCKAP1 expression

To identify the responsible targets related to ALS(R)-iMGs phagocytic dysfunction, transcriptomic data were analyzed in another set of iMGs from ALS(R) (R6 – R8) and ALS(S) (S7 – S10) patients. In principle component analysis (PCA), ALS(R)-iMGs were distinct from ALS(S)-iMGs (Figure 2A), and DEG analysis (fold changes > 1.5) showed that 2,599 genes were differentially expressed in ALS(R)-iMGs (Figure 2B and Supplemental Table 6). Gene ontology (GO) analysis of 2599 genes revealed that gene subsets, including response to lipopolysaccharide, inflammatory response, actin filament polymerization, and phagocytosis, were differentially expressed between ALS(R)-iMGs and ALS(S)-iMGs (Figure 2C and Supplemental Table 7).

**Figure 2:**
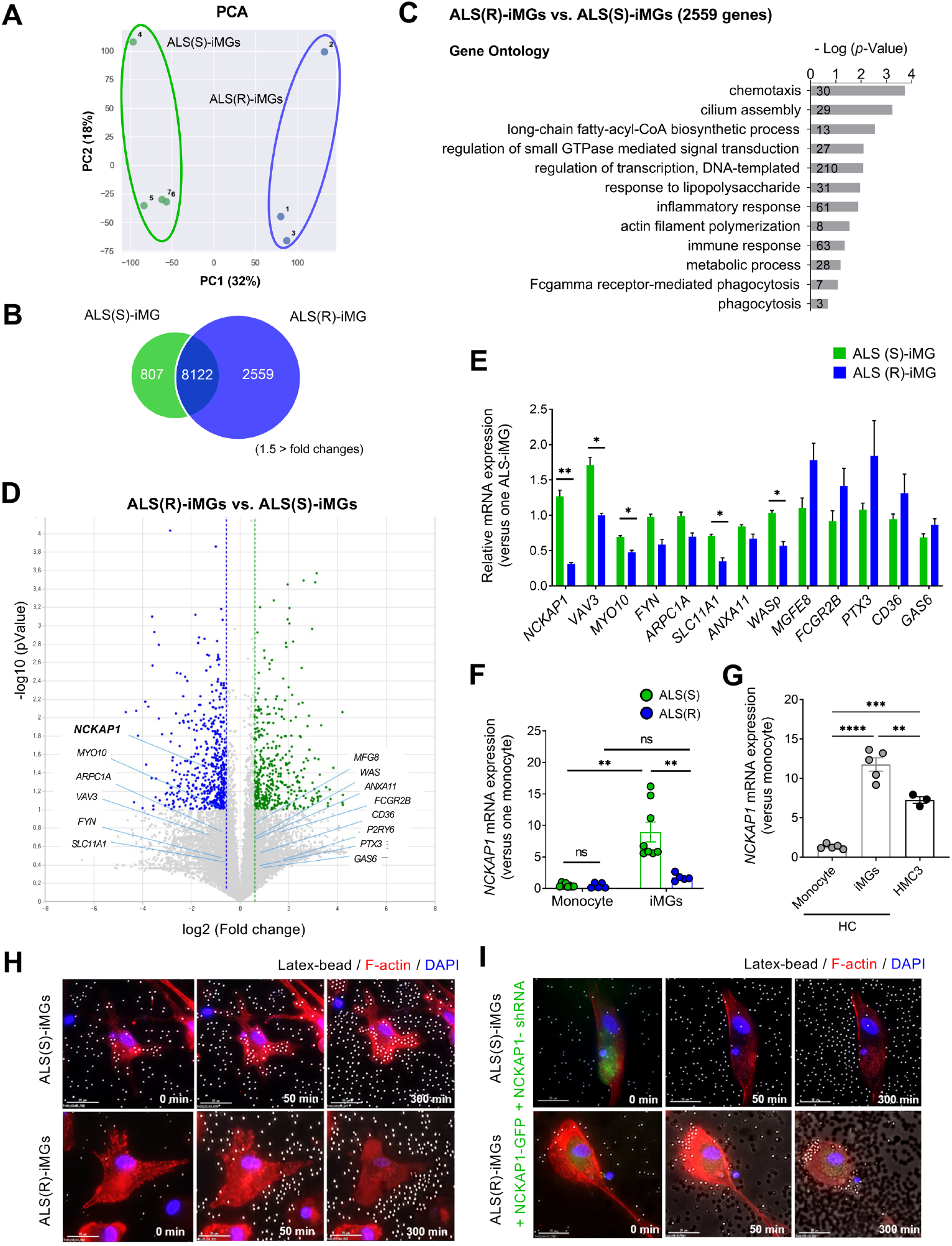
*NCKAP1* is associated with impaired phagocytic function in ALS(R)-iMGs. Transcriptome analysis of iMGs generated from three ALS(R) and four ALS(S) patients. **(A)** Principal component analysis (PCA) of all expressed genes in individual ALS(R)-iMGs and ALS(S)-iMGs. **(B)** Venn diagram illustrating that the number and overlap of transcripts differed significantly in the two groups according to RNA-seq (1.5-fold-change). **(C)** Functional analysis using DAVID software from 2,559 transcripts that were significantly altered in ALS(R)-iMGs compared to ALS(S)-iMGs (10 GO analyses). The number within each bar indicates the number of genes in the database for the specified term. **(D)** Volcano plot showing transcripts with significantly altered abundance in ALS(R)-iMGs compared to ALS(S)-iMGs. We tested for significant differences between ALS(R)-iMGs and ALS(S)-iMGs samples, which are highlighted, using a cutoff of *P*-values of 0.05 with a 1.5-fold-change ratio cutoff. GO genes of phagocytosis (GO: 0006909) are presented. **(E)** A subset of 13 transcripts identified as ‘hits’ (phagocytosis-related increased or decreased abundance) in ALS(R)-iMGs compared to ALS(S)-iMGs were validated by RT-qPCR. **(F)** *NCKAP1* mRNA expression in individual-subject-derived monocytes or iMGs from the two groups. The symbols represent individual-subject-derived monocytes or iMGs (ALS(R): n = 5; ALS(S): n = 8, from available samples). **(G)** *NCKAP1* mRNA expression in HC-iMG and human microglial cell (HMC3) compared to HC-monocytes (n = 5). **(H)** Snapshots of live-cell imaging showing the phagocytosis of latex beads in both groups of ALS-iMGs. Merged images of DIC (latex bead) and fluorescence images (F-actin: red; DAPI: blue). ALS(S)-iMGs (upper) showed normal phagocytic function, whereas ALS(R)-iMGs (lower) showed a marked impairment in phagocytosis. Representative frames from a time-lapse image series (0-5 h) are shown. The figure represents independent experiments performed in replicates (n = 10, ALS(R)-iMGs: n = 7; ALS(S)-iMGs: n = 4). **(I)** Snapshots of live-cell imaging showing the phagocytosis of latex beads in ALS(S)-iMGs transfected with shNCKAP1-GFP (upper) or ALS(R)-iMGs transfected with NCKAP1-GFP (lower). The impaired phagocytic function in ALS(R)-iMGs was rescued by NCKAP1 overexpression. Representative frames from a time-lapse image series (0 5 h) are shown. The figure represents independent experiments performed in replicates (n = 10). Data were first tested for normality test. Statistical analysis between two groups was performed using Unpaired t-test (E), Mann Whitney test (F), or Wilcoxon matched-pairs signed rank test (F; monocyte vs. iMGs) and between multiple groups using one-way ANOVA with Tukey analysis (G). Values are the mean ± SEM (*P < 0.05, **P < 0.01, ***P < 0.001, ****P < 0.0001, ns: not significant). Scale bar: 25 μm.

NCK-associated protein 1 (NCKAP1) was the first candidate identified related to defective phagocytosis (fold changes > 1.5) in ALS(R)-iMGs (Figure 2D). Thirteen candidate transcripts were investigated in all available samples by RT-qPCR. Five genes (*NCKAP1, VAV3, MYO10, SLC11A1*, and *WASp*) reproduced the RNA-Seq results. *NCKAP1*, guanine nucleotide exchange factor Vav 3 (*VAV3*), *MYO10*, and Wiskott-Aldrich syndrome protein (*WASp*), known intracellular signaling-regulated factors related to the actin-polymerization process in phagocytosis (21), were significantly decreased in ALS(R)-iMGs. The mRNA expression level of *NCKAP1* in individual monocytes was barely detectable, but iMGs of both groups expressed higher *NCKAP1* than the monocytes (Figure 2F). HC-iMGs and human microglial cell (HMC3) also showed higher *NCKAP1* expression than monocytes (Figure 2G).

To further delineate the role of NCKAP1 in phagocytosis, we generated another set of iMGs from ALS(R) (R9 – R15) and ALS(S) (S11 – S14) patients. NCKAP1 overexpression rescued the defective phagocytic function of ALS(R)-iMGs using live-cell imaging. While ALS(S)-iMGs phagocytized latex beads (Figure 2H, upper panel), ALS(R)-iMGs showed defective phagocytosis (Figure 2H, lower panel). However, when NCKAP1 was overexpressed in ALS(R)-iMGs, phagocytosis was rescued (Figure 2I, lower panel). As expected, the active phagocytosis in ALS(S)-iMGs was not present when NCKAP1 was knocked down (Figure 2I, upper panel).

### miRNA-214-3p and miRNA-34-3p targeting NCKAP1 are associated with NCKAP1 reduction and progression speed in the discovery cohort

To leverage our findings to serological biological markers, we searched miRNAs targeting NCKAP1. miRNA targets were predicted by combinatorial utilization of the TargetScan and miRDB web-based prediction algorithms, and the collected targets were experimentally confirmed *in vitro* and in ALS-related and inflammation-related miRNAs from previous studies (22-26). We identified putative miR-214-3p and −34c-3p binding sites in the 3′-UTR of NCKAP1 using bioinformatics tools for miRNA target prediction. To confirm, we tested the effects of each mimic and inhibitor in HeLa cells. Both miR-214-3p and −34c-3p directly downregulated NCKAP1, whereas the inhibitors upregulated NCKAP1 (Figure 3A) (22, 23). These findings indicated that both miR-214-3p and miR-34c-3p played a role in regulating NCKAP1 levels.

**Figure 3:**
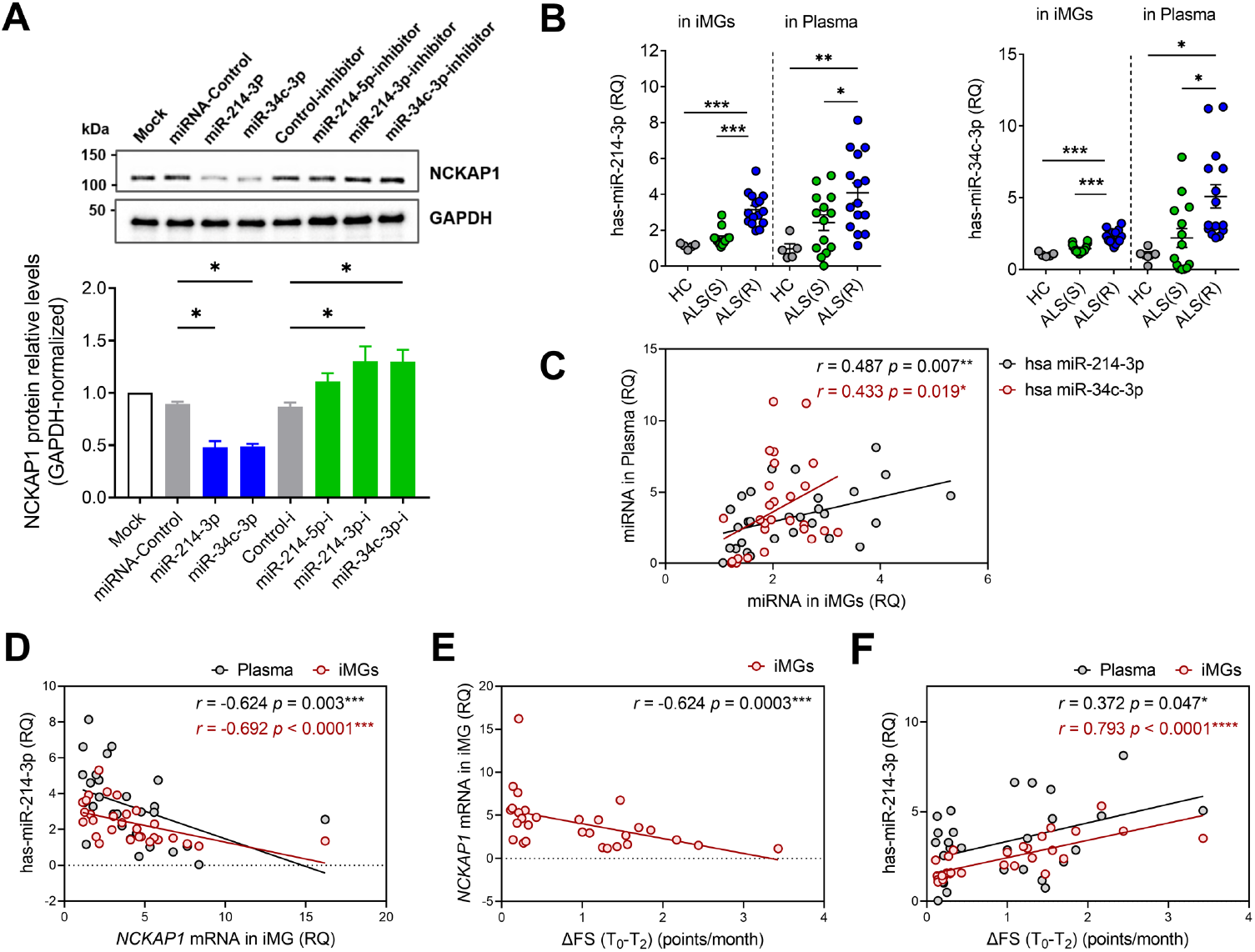
*miR-214 and miR-34c* as candidate biomarkers targeting NCKAP1 and predicting the progression speed of ALS. **(A)** Immunoblot for NCKAP1 expression in HeLa cells transfected with miR-214-3p/miR-34c-3p mimic, inhibitor, or negative control. Quantification of NCKAP1 protein level was performed via normalization to GAPDH (n = 3). **(B)** Relative expression of miR-214-3p and miR-34c-3p in plasma and iMGs of discovery cohort. Each data point represents one participant (HC: n = 5; ALS(R): n = 15; ALS(S): n = 14). **(C)** Correlation of the relative expression of miR-214-3p and miR-34c-3p between plasma and iMGs. **(D)** Correlation of the relative expression of miR-214-3p in plasma or iMGs with the relative expression of *NCKAP1* in iMGs. **(E)** Correlation of the relative expression of *NCKAP1* in iMGs with the rate of disease progression from the symptom onset time (T_0_) to the sampling time point for the generation of iMGs (T_2_) (ΔFS, points/month). **(F)** Correlation of the relative expression of miR-214-3p in plasma or iMGs with disease progression. Each data point in the correlation study represents one participant (ALS(R): n = 15; ALS(S): n = 14). Data were first tested for normality test. Statistical analysis between multiple groups was performed using one-way ANOVA with Tukey analysis (A, B) or Kruskal-Wallis with post-hoc Dunn’s analysis (B). Correlations were analyzed by Spearman rank correlation test (C∼F). Values are the mean ± SEM (*P < 0.05, **P < 0.01, ***P < 0.001, ****P < 0.0001).

Based on the experimental results, we selected NCKAP1 and its regulators (miR-214-3p and −34c-3p) as candidate biomarkers reflecting the phagocytic function of ALS-iMGs and the clinical progression of ALS. In the discovery cohort, an internal validation study was performed to evaluate whether ΔFS (T_0_-T_2_) was correlated with the candidate miRNAs. First, we confirmed that each miRNA in both iMGs and plasma samples was elevated in ALS(R) compared to ALS(s) (Figure 3B). miRNA levels directly measured in iMGs were correlated with plasma miRNA (miR-214-3p: *p* = 0.007; miR-34c-3p: *p* = 0.019; Figure 3C). Each miRNA in iMGs and plasma was negatively correlated with *NCKAP1* mRNA levels (plasma: *p* = 0.003; iMGs: *p* < 0.0001; Figure 3D). The final step of internal validation was an analysis of the relationship between patient progression speed and *NCKAP1* mRNA, and candidate miRNA biomarkers. ΔFS (T_0_-T_2_) was negatively correlated with iMGs *NCKAP1* mRNA levels (*r = -*0.624, *p* = 0.0003; Figure 3E). ΔFS (T_0_-T_2_) was also correlated with miR-214-3p in both iMGs and plasma sampled at T_2_ (iMGs: *r=*0.793, *p* < 0.0001, plasma: *r =* 0.372, *p* = 0.047; Figure 3F). Based on the serial data of the discovery cohort, we selected plasma miR-214-3p as a candidate biomarker to reflect the progression speed of ALS from symptom onset to sampling time.

### Plasma miRNA-214-3p as a reliable biomarker predicting ALS progression speed

To validate the reliability of the discovered biomarker, we evaluated whether plasma miR-214-3p could be a serological biomarker predicting the progression speed of ALS patients in the validation cohort. Serial ALSFRS-R data recorded at T_1_ and T_2_ and bio-banked frozen plasma collected at T_1_ were used for validating miR-214-3p with ΔFS (T_1_-T_2_). In this cohort, the participants were randomly sub-grouped into quadruplets (Q1, Q2, Q3, Q4) according to ΔFS (T_1_-T_2_) data based on ALSFRS-R measured at T_1_ and T_2_. We classified the enrolled patients into three groups according to ΔFS as rapidly progressive ALS [ALS(R)] (Q1, ΔFS (T_1_-T_2_) > 1.45), intermediate ALS [ALS(I)] (Q2 and Q3, 1.45 ≥ ΔFS (T_1_-T_2_) ≥ 0.46), and slowly progressive ALS [ALS(S)] (Q4, ΔFS (T_1_-T_2_) < 0.46) (Table 2 and Supplemental Table 2).

**Table 2.** Demographics and characteristics of the validation cohort.

| Characteristic | Rapid-ALS (Q1) | Intermediate-ALS (Q2, Q3) | Slow-ALS (Q4) | Total-ALS | Controls | *p* |
| --- | --- | --- | --- | --- | --- | --- |
| Number of subjects | 32 | 67 | 33 | 132 | 30 | — |
| Male: Female, n | 21:11 | 39:28 | 16:17 | 76:56 | 15:15 | — |
| Age at symptom onset (T<sub>0</sub>), year | 60.7 (8.9) | 57.9 (11.0) | 56.6 (10.9) | 58.2 (10.5) | — | $p^e = 0.237$<br>$p^f = 0.430$<br>$p^g = 0.826$ |
| Age at the baseline point at initial visit time for diagnosis (T<sub>1</sub>), year | 61.6 (9.0) | 59.3 (10.9) | 59.0 (11.1) | 59.8 (10.5) | 59.6 (6.4) | $p^e > 0.999$<br>$p^f > 0.999$<br>$p^g > 0.999$ |
| ALSFRS-R recorded at T<sub>1</sub> | 36.3 (5.6) | 38.7 (5.3) | 41.2 (4.5) | 38.8 (5.4) | — | $p^e < 0.001^*$<br>$p^f = 0.088$<br>$p^g = 0.057$ |
| Disease duration (T<sub>0</sub>-T<sub>1</sub>)<sup>a</sup>, months | 10.3 (8.0) | 16.4 (14.0) | 28.4 (33.2) | 17.9 (20.6) | — | $p^e < 0.001^*$<br>$p^f = 0.008^*$<br>$p^g = 0.172$ |
| ΔFS (T<sub>0</sub>-T<sub>1</sub>)<sup>b</sup>, point/months | 1.4 (0.8) | 0.8 (0.5) | 0.4 (0.2) | 0.8 (0.7) | — | $p^e < 0.001^*$<br>$p^f < 0.001^*$<br>$p^g < 0.001^*$ |
| Age at the time point for recording ALSFRS-R, at least six months the baseline point (T<sub>2</sub>), year | 62.2 (9.0) | 60.0 (10.8) | 59.6 (11.2) | 60.4 (10.5) | — | $p^e > 0.999$<br>$p^f > 0.999$<br>$p^g > 0.999$ |
| **ALSFRS-R measured at T<sub>2</sub>** | 19.1 (9.4) | 31.5 (6.3) | 39.4 (4.5) | 30.5 (9.9) | — | $p^e < 0.001^*$<br>$p^f < 0.001^*$<br>$p^g < 0.001^*$ |
| **Disease duration (T<sub>1</sub>-T<sub>2</sub>)<sup>c</sup>, months** | 6.7 (1.1) | 7.9 (2.8) | 7.4 (1.7) | 7.5 (2.3) | — | $p^e = 0.173$<br>$p^f = 0.109$<br>$p^g > 0.999$ |
| **ΔFS (T<sub>1</sub>-T<sub>2</sub>)<sup>d</sup>, point/months** | 2.6 (1.0) | 0.9 (0.3) | 0.2 (0.1) | 1.2 (1.0) | — | $p^e < 0.001^*$<br>$p^f < 0.001^*$<br>$p^g < 0.001^*$ |
| **Family history of ALS, n (%)** | 0 (0%) | 0 (0%) | 0 (0%) | 0 (0%) | 0 (0%) | — |

In the validation cohort, the symptom onset time was defined as T<sub>0</sub>, the baseline point at the initial visit time for diagnosis or treatment as T<sub>1</sub>, and the time point for recording ALSFRS-R, at least six months after the baseline point (T<sub>1</sub>) as T<sub>2</sub>. We used the bio-banked sample collected at the initial visit time (T<sub>1</sub>) for miRNA validation analysis.
Data are means (standard deviation) for continuous variables or n (%) for categorical variables.
<sup>a</sup>Disease duration: the period from symptom onset time (T<sub>0</sub>) to the baseline point at the initial visit time (T<sub>1</sub>).
<sup>b</sup>The progression rate was defined as ΔFS (T<sub>0</sub>-T<sub>1</sub>), (i.e., (48 - ALSFRS-R score at T<sub>1</sub>) / (duration from T<sub>0</sub> to T<sub>1</sub>, months)).
<sup>c</sup>Disease duration: the period from the baseline point at the initial visit time (T<sub>1</sub>) to the time point for recording ALSFRS-R, at least six months after the baseline point (T<sub>2</sub>).
<sup>d</sup>The progression rate was defined as ΔFS (T<sub>1</sub>-T<sub>2</sub>), (i.e., (ALSFRS-R at T<sub>1</sub>) - (ALSFRS-R at T<sub>2</sub>) / (duration from T<sub>1</sub> to T<sub>2</sub>, months)). ALS patients were categorized as rapidly progressive ( ΔFS (T<sub>1</sub>-T<sub>2</sub>) > 1.45), intermediate ALS ( $0.46 \leq \Delta$ FS (T<sub>1</sub>-T<sub>2</sub>) $\leq 1.45$ ), and slowly progressive ( ΔFS (T<sub>1</sub>-T<sub>2</sub>) < 0.46).
Statistical analysis was performed using an unpaired t-test or one-way ANOVA (to compare multiple treatment groups versus control) with $*p < 0.05$ indicating a significant difference.
<sup>e</sup> $p$ -value: comparison between rapid-ALS group (Q1) and slow-ALS groups (Q4).
<sup>f</sup> $p$ -value: comparison between rapid-ALS (Q1) and intermediate-ALS groups (Q2, Q3).

Plasma miR-214-3p was significantly higher in patients with ALS than in HCs (Figure 4A), and ROC curves indicated that plasma miR-214-3p was a potential diagnostic biomarker for ALS, with AUC values of 0.839 (*p* < 0.0001; Figure 4B). Plasma miR-214-3p was increased in ALS(R), ALS(I), and ALS(S) compared to HCs (Figure 4C). ROC curves for miR-214-3p showed an AUC of 0.657 for discrimination between ALS(R) and ALS(S) (Figure 4D, *p* = 0.029). Finally, plasma miR-214-3p levels were positively correlated with disease progression speeds, at both ΔFS (T_0_-T_1_) (*r* =0.297, *p* = 0.0005) and ΔFS (T_1_-T_2_) (*r* =0.230, *p* = 0.0079) (Figure 4E).

**Figure 4:**
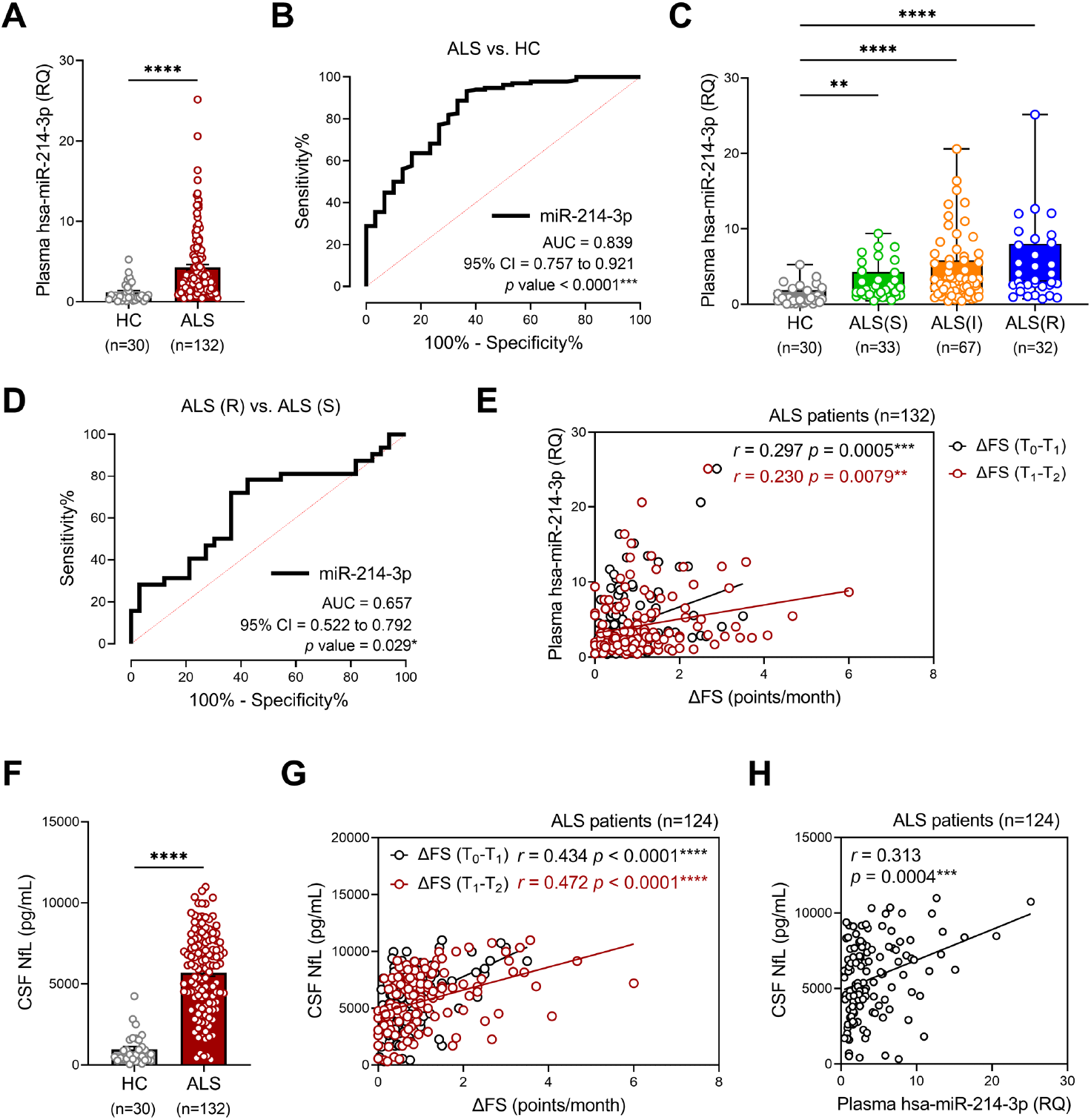
miRNA-214 as a predicting biomarker of ALS progression speed in validation cohort. Plasma miRNA-214-3p level was measured in the validation cohort (132 ALS patients and 30 HCs). The log-normalized relative abundance of miRNA-214-3p between two groups. ROC curve analyses of miRNA-214-3p in ALS vs. HC. AUC, 95% CI, and *P*-values are shown. **(C)** Plasma miRNA-214-3p levels in ALS(R), ALS(I), and ALS(S) (by ΔFS from the baseline point at the initial visit time for diagnosis (T_1_) to the time point for obtaining ALSFRS-R, at least six months after the initial visit (T_2_)) and HCs. **(D)** ROC curve analysis for discrimination between ALS(R) and ALS(S). AUC, 95% CI, and *P*-values are shown. **(E)** Correlation of miR-214-3p in plasma with the rate of disease progression (black line; ΔFS from symptom onset time (T_0_) to the baseline point at the initial visit time for diagnosis (T_1_), red line; ΔFS from the sampling time at the baseline point at the initial visit time (T_1_) to the time point for obtaining ALSFRS-R, at least six months after the baseline point (T_2_), points/month). **(F)** CSF NfL levels in the validation cohort (124 ALS patients and 30 HCs). **(G)** Correlation of NfL in CSF with the rate of disease progression (black line; ΔFS from T_0_ to T_1_, red line; ΔFS from T_1_ to T_2_, points/month). **(H)** Correlation of plasma miR-214-3p with CSF NfL. Each data point represents one participant. Data were first tested for normality test. Statistical analysis between two groups was performed using Mann Whitney test (A, F) and between multiple groups was performed using Kruskal-Wallis with post-hoc Dunn’s analysis (C). Correlations were analyzed by Spearman rank correlation test (E, G, and H). Values are the mean ± SEM (*P < 0.05, **P < 0.01, ***P < 0.001, ****P < 0.0001).

NfLs are promising candidate biomarkers with prognostic implications in ALS, and CSF NfL concentrations correlated with survival and disease progression (2, 27, 28). We assayed NfLs using CSF samples bio-banked at T_1_ (124 ALS, 30 HCs). CSF NfLs were elevated in ALS (mean, 5697 pg/mL) compared to HCs (960.6 pg/mL, *p* < 0.001; Figure 4F) and correlated with disease progression speeds at both ΔFS (T_0_-T_1_) (*r* =0.434, *p* < 0.0001) and ΔFS (T_1_-T_2_) (*r* =0.472, *p* < 0.0001; Figure 4G). Plasma miR-214-3p was correlated with CSF NfLs (*r* = 0.313, *p* = 0.0004; Figure 4H). Therefore, miR-214-3p could be a candidate biomarker for predicting future progression speed related to phagocytic dysfunction in ALS patients.

## DISCUSSION

This study identified possible therapeutic targets in rapidly progressive ALS and new biological markers predicting progression speed related to phagocytic dysfunction using iMGs model. Only ALS(R)-iMGs showed intrinsically perturbed phagocytosis. Using transcriptomic analysis, we identified defective phagocytosis corresponding to reduced NCKAP1 levels as the key factor promoting phagocytic dysfunction in ALS(R)-iMGs, suggesting that NCKAP1 reduction in ALS(R)-iMGs may be responsible for defective phagocytosis. miR-214-3p and miR-34c-3p were identified as negative regulators of NCKAP1. In the discovery cohort, miR-214-3p and miR-34c-3p levels were correlated with NCKAP1 and the ΔFS of ALS patients. miR-214-3p was selected from the internal validation data as a candidate biomarker predicting ALS progression speed. The validation study conducted in 132 ALS patients and 30 HCs confirmed that plasma miR-214-3p could be used as a new candidate biomarker to predict the progression speed reflecting phagocytic dysfunction of ALS patients.

Only a few studies have reported an association between NCKAP1 and neurodegenerative diseases (29). Most studies on NCKAP1 have focused on its role in neuronal differentiation and migration as a cytoskeletal regulator during development (30). *NCKAP1L*, a hematopoietic cell-specific gene with a structure similar to *NCKAP1*, was reported as a key player in actin polymerization and essential for macrophage migration and phagocytosis (31, 32). In contrast to NCKAP1L, NCKAP1 is enriched in the brain but absent or less expressed in hematopoietic cells (33). Given that NCKAP1 and NCKAP1L are members of the WAVE2 regulatory complex (WRC) that can promote actin polymerization, low NCKAP1 expression may cause complex instability and hinder F-actin polymerization in iMGs. Therefore, NCKAP1 is presumed to play a key role in phagocytosis engulfment by regulating actin cytoskeleton dynamics.

Biological markers such as NfL and TDP-43 currently used to predict the likelihood of rapid disease progression in ALS patients have limited sensitivity due to clinical and genetic heterogeneity (2, 27). And, CSF NfL measurements require a relatively invasive procedure. Therefore, a reliable and easily accessible biomarker compatible with CSF NfLs is needed. In this study, we found that NCKAP1 reduction was associated with miR-214-3p increase in ALS(R)-iMGs. In addition, plasma miR-213-3p showed positive correlation with miR-213-3p level in ALS-iMGs in discovery cohort. Given that measuring NCKAP1 and miR-214-3p in iMGs is expensive and time-consuming, plasma miR-213-3p can be a simple and easily accessible serological biomarkers because miRNAs have recently attracted attention as potential biomarkers for ALS diagnostic or prognostic purposes due to their easy detection and stable status in blood and body fluids (34). In addition, miR-214-3p expression was increased in microglia isolated from *SOD1*-G93A mice (24), and the muscles (25) and sera (26) of ALS patients. Collectively, our study proposes a new plasma biomarker reflecting phagocytic dysfunction and progression speed in ALS patients.

The validation cohort was designed to validate whether the discovered candidate could predict future ALS progression speed. We analyzed miR-214-3p using blood samples bio-banked at the initial visit time (T_1_) in patients who were followed for more than six months from the baseline point. We conducted a validation study of prognostic biomarkers through correlation analysis of participant disease progression speed between T_1_ and the follow-up time (T_2_). And we evaluated whether miR-214-3p reflected past progression speed from onset (T_0_) to T_1_ using plasma bio-banked at T_1_. Plasma miR-214-3p was correlated with ΔFS (T_0_-T_1_) (*r* =0.297, *p* = 0.0005), ΔFS (T_1_-T_2_) (*r* =0.230, *p* = 0.0079), ΔFS (T_0_-T_2_) (*r* =0.285, *p* = 0.0009, data not shown), and NfL (*r* =0.313, *p* = 0.00.4). The validation study showed that plasma miR-214-3p was closely correlated with patient ΔFS, reflecting ALS disease progression speed. Plasma miRNA was measured in bio-banked T_1_ samples, and miRNA levels were validated to determine whether it had predictive power for progression speed over at least the next six months. This data implied that miRNA levels measured at the initial visit for diagnosis or therapeutic intervention could be used as a reliable biomarker to predict future patient progression speed or prognosis. However, the internal standardization of miRNA measurements and age-related and other factor-related confounding effects should be considered before its practical application. These findings suggest that the T_1_ concentrations of miR-214-3p could provide information on anticipated progression speed, allowing the stratification of ALS patients in clinical trials.

CSF NfL concentrations in ALS patients reportedly correlated with both survival and the disease progression rate (2, 27, 28). Therefore, we compared the correlative power of miR-214-3p and CSF NfL, the best current biomarker to predict prognosis, with ΔFS. In the current study, the correlative value of miR-214 with ΔFS was significant (*r* = 0.256), but not inferior to the baseline CSF NfL (*r* = 0.417) predicting the anticipated ΔFS in the validation cohort. Nevertheless, miR-214-3p has several merits as a promising biomarker. First, it can be easily measured from plasma and serially monitored as a scalable unit. Second, plasma miR-214-3pcould be used as a unique potential biomarker to verify immune-inflammatory target engagement and monitor the intended therapeutic efficacy in therapeutic strategies focused on the functional modulation of microglia.

However, this study had limitations. First, we only focused on the relationship between miR-214-3p, disease progression speed, and iMGs phagocytic function. Considering that miRNA has diverse targets, the exact role of miR-214-3p has not yet been entirely revealed. Therefore, further studies, including the temporal profile of plasma miR-214-3p in longitudinal studies and comparative analysis with other biomarkers in a larger cohort should be conducted to confirm miR-214-3p as a biomarker predicting the anticipated progression of ALS. The current study excluded patients with familial or causative genetic phenotypes, and frontotemporal dementia, and only included sporadic ALS (Supplemental Table 1 and 2). Therefore, we could exclude these confounding or heterogeneous clinical-genetic factors, which might influence miR-214-3p concentrations. Second, we used the same internal control to standardize the miR-214-3p quantification according to the manufacturer’s guidelines for profiling biofluid miRNAs. Therefore, the development of accurate standardization of miR-214-3p measurements is required to avoid interlaboratory variations and for use in future clinical trials for the stratification of patients.

We discovered the potential of miR-214 as a novel candidate biomarker predicting progression speed. miR-214-3p measurements allow the stratification of ALS patients with rapidly or slowly progressive disease. Moreover, patient prognosis based on the miR-214-3p biomarker could help in the development of tailored, targeted therapies to prolong or improve ALS patients’ quality of life. In conclusion, our study proposed a new candidate biomarker, miRNA-214-3p, to predict ALS progression speed and for use as a possible therapeutic target to rescue microglial phagocytic dysfunction in rapidly progressive ALS patients.

## METHODS

### Study design and sample preparation

We designed the current study as a biomarker discovery study (discovery cohort)—the discovery of candidate biomarkers related to phagocytic dysfunction and ALS progression speed using a monocyte-derived iMGs model—and a validation study of the discovered candidate for its clinical usability in predicting disease progression speed (validation cohort).

All participants had clinically definite, clinically probable, or clinically probable with a laboratory-supported diagnosis according to the revised El Escorial criteria (35). The patients’ clinical information, including ALS Functional Rating Scale–Revised (ALSFRS-R) score (0∼48), was prospectively registered in the database. Blood and cerebrospinal fluid (CSF) samples had been banked since the initial visit (9). We defined the symptom onset time as T_0_ and the baseline point at the initial visit for diagnosis or treatment as T_1_. T_2_ was defined as the sampling time point for the generation of iMGs in the discovery cohort. T_2_ was the time point for regular ALSFRS-R recording in the validation cohort at least six months after T_1_. Progression speed was expressed as delta FS (ΔFS) (36). In general, the ΔFS between the first time point (A) and the second time point (B) was calculated as ΔFS = [(ALSFRS-R at A) – (ALSFRS-R at B) / (duration from B to A, months)]. However, we used a modified variable calculated at the time of sampling or interest into the study as: ΔFS (T_0_-T_1_) between T_0_ and T_1_, ΔFS (T_1_-T_2_) between T_1_ and T_2_, and ΔFS (T_0_-T_2_) between T_0_ and T_2_.

We enrolled 29 participants with ALS and five healthy volunteers in the biomarker discovery study between September 2015 and July 2017. We tentatively sub-grouped participants who exhibited extremely rapid or slow progression in the ALS/MND Clinic database at Hanyang University Hospital as having rapidly progressive ALS [ALS(R); ΔFS (T_0_-T_2_) ≥ 1.0, n=15] and slowly progressive ALS [ALS(S); ΔFS (T_0_-T_2_) < 0.5, n=14] (28, 37). After obtaining informed consent, blood samples were collected to generate iMGs, and comparative studies to discover candidate biomarkers were serially conducted. All participants’ medical records were reviewed, and their clinical characteristics are summarized in Table 1 and Supplemental Table 1.

In the validation study, we enrolled 132 participants with ALS who were followed more than six months since T_1_ and 30 HCs. Patient clinical characteristics are summarized in Table 2 and Supplemental Table 2. We excluded participants with familial history or pathogenic variants of ALS-related genes in whole-exome sequencing to exclude possible genetic effects on ALS progression (Supplemental Table 3). To verify our data on progression speed and candidate biomarkers, we analyzed plasma inflammatory cytokines in the discovery cohort and neurofilaments in the validation cohort, respectively. Plasma interleukin (IL)-6, interferon (IFN)-γ, IL-8, tumor necrosis factor (TNF)-α, and monocyte chemoattractant protein (MCP)-1 were measured with ELISA kits (R&D Systems, Minneapolis, MN, USA), according to the manufacturer’s instructions, and CSF NfLs were measured with an ELISA kit (UmanDiagnostics AB, Umeå, Sweden). HC samples were collected from ALS patient spouses after obtaining consent.

### Establishment of induced microglia-like cells (iMGs) from PBMCs

PBMCs were isolated by density gradient centrifugation using Ficoll (GE Healthcare, Uppsala, Sweden). iMG cells were established using a previously published method (17). Briefly, PBMCs were resuspended in RPMI-1640 (Gibco, Carlsbad, CA, USA) containing 10% fetal bovine serum (FBS; Gibco) and 1% antibiotic/antimycotic (Invitrogen, Carlsbad, CA, USA) and cultured overnight at 37 °C and 5% CO2. The next day, adherent cells (monocytes) were cultured in RPMI-1640 Glutamax (Gibco) supplemented with 1% antibiotic/antimycotic, recombinant granulocyte-macrophage colony-stimulating factor (GM-CSF) (R&D Systems), and recombinant IL-34 (IL-34) (R&D Systems) to develop iMG cells. After generating iMGs, the cells were labeled with human CD11b-APCVio770 and CD45-phycoerythrin (PE) (Miltenyi Biotec, Gladbach, Germany), and flow cytometry was performed as described previously (38). All data were assessed by FACSCanto II flow cytometry (BD Biosciences, Piscataway, NJ, USA) and analyzed by FlowJo software. Gene expression in iMGs was measured using quantitative real-time polymerase chain reaction (RT-qPCR) as described previously (38). Primer information is shown in Supplemental Table 4.

### Immunostaining and morphology analysis

To visualize iMGs, the cells were fixed and stained with a microglial marker (IBA1) and counterstained with DAPI (4′,6-diamidino-2-phenylindole). Immunostaining was performed as previously described (38). Antibody information is shown in Supplemental Table 4. Images were acquired by confocal microscopy (TCS SP5, Leica, Wetzlar, Germany). Three-dimensional reconstructions of randomly selected iMG cells (IBA1-positive) were generated using Imaris software (Bitplane, Zurich, Switzerland). Two blinded researchers performed morphometric analysis of each reconstructed cell after determining dendrite length, the number of segments, and branch points (39).

### Phagocytosis assay

iMGs were treated with 4 µl of red fluorescent latex beads for 24 h at 37 °C. The cells were washed twice with ice-cold phosphate-buffered saline (PBS), fixed, and stained with a microglial marker (P2RY12) and DAPI. The number of phagocytized beads was counted using ImageJ software (40). To assess phagocytosis cup formation in iMGs, cells were incubated with latex beads for 2 h, fixed, and stained with fluorescent phalloidin (1: 1,000; Molecular Probes, Eugene, OR, USA) for 45 min with secondary antibodies. Antibody information is shown in Supplemental Table 4. Images were acquired by confocal microscopy. For live-cell imaging, iMGs were grown in Lab-Tek II Chamber Slide (Thermo Fisher Scientific, Rochester, NY, USA) and labeled with 100 nM SiR-actin dye (Cytoskeleton Inc., Denver, CO, USA) according to the manufacturer’s protocol. Beads (1.1 μm, Sigma-Aldrich, Saint Louis, MO, USA) were added to the cells before analysis. Images were captured one frame every 1 min 30 sec over 5 h using a DeltaVision Imaging System (Applied Precision, Bratislava, Slovakia).

### RNA sequencing and data analysis

RNA quality was assessed with an Agilent 2100 bioanalyzer using an RNA 6000 Nano Chip (Agilent Technologies, Amstelveen, Netherlands). RNA libraries were constructed using a SENSE 3′ mRNA-Seq Library Prep Kit (Lexogen, Inc., Vienna, Austria) according to the manufacturer’s instructions. High-throughput sequencing was performed as single-end 75 sequencings using a NextSeq 500 platform (Illumina, Inc., San Diego, CA, USA). SENSE 3′ mRNA-Seq reads were aligned using Bowtie2 version 2.1.0 (41). Differentially expressed genes (DEGs) were determined based on counts from unique and multiple alignments using EdgeR in R version 3.2.2 and BIOCONDUCTOR version 3.0 (42). Read count data were processed based on the global normalization method using Genowiz™ version 4.0.5.6 (Ocimum Biosolutions, India). Gene classification was based on DAVID (http://david.abcc.ncifcrf.gov/)database searches. MeV 4.9.0 was used for clustering samples and genes and visualization.

### Cell culture and transfection

Human microglial clone 3 (HMC3) cells (ATCC, Manassas, VA, USA) were cultured in Eagle’s Minimum Essential Medium (MEM) containing 10% FBS and antibiotics. HeLa cells were cultured in Dulbecco’s MEM containing 10% FBS, sodium bicarbonate, sodium pyruvate (Sigma-Aldrich), and antibiotics. HeLa cells were transfected with green fluorescent protein (GFP)-tagged human NCKAP1 cDNA or NCKAP1 shRNA using Lipofectamine 2000 (Invitrogen) according to the manufacturer’s protocol. iMGs were transduced with pLenti-C-mGFP-Human NCKAP1 (NM_013436), the cDNA ORF Clone (OriGene Technologies, Rockville, MD, USA), or pGFP-C-shLenti-NCKAP1 Human shRNA lentiviral particles (ID 10787) according to the manufacturer’s protocol. For miRNA modulation, miRNA mimics or inhibitors and control oligonucleotides were synthesized by Bioneer Corporation (Daejeon, Korea) and transfected using RNAiMAX (Invitrogen) into HeLa cells following the manufacturer’s protocol. After transfection, cells were assessed by western blotting and RT-qPCR as previously described (38). Antibody, primers, and miRNA information are shown in Supplemental Table 4.

### miRNA analysis

miRNAs of iMGs were isolated using the miRNeasy Mini Kit (Qiagen, Hilden, Germany), reverse transcribed using the miScript II RT Kit (Qiagen), and cDNA was diluted 1:50. Plasma miRNAs were isolated using the miRNeasy Serum/Plasma Advanced Kit (Qiagen) and retrotranscribed by the miRCURY LNA RT kit (Qiagen) including spike-in cel-miR-39. cDNA (250 ng) was used for RT-qPCR in the Step One Plus system (Applied Biosystems), using miRCURY LNA SYBR Green PCR Master Mix and miRCURY LNA miRNA PCR primers (Qiagen). All protocols were performed according to the manufacturers’ instructions. After assessing threshold cycle (Ct) values, the relative expression levels of miRNA were calculated by the ΔΔCt method using U6 small RNA (in iMGs). miR-103-3p (in plasma) served as the normalization control.

### Sample size calculation

To validate the difference in miR-214 expression between rapid and slow progressors, we collected 132 samples based on the sample size calculation using R pwr packages (pwr.t.test). Effect size (Cohen’s d) was calculated by Eq (1). Pooled standard deviation (SD_pooled_) was calculated by Eq (2).

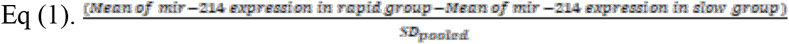

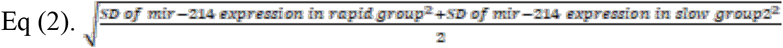

We set an alpha(α) of 0.05, and a power(1-β) of 0.2. Based on the study set, the pooled standard deviation and effect size were 1.058 and 0.487, respectively. Using the function pwr.t.test (d = 2.192, sig.level = 0.05, power = 0.8, type = “two.sample”, alternative = “two.sided”) and rounding up to an integer, we obtained five samples per group.

### Statistics

Data are presented as either mean ± SD or SEM. Normality was accessed by Shapiro–Wilk and Kolmogorov-Smirnov normality tests. For normal distribution, comparisons between ALS and HC cohorts were performed using the Unpaired t test or Paired t test (two-tailed) while multiple comparisons were performed using ordinary one-way ANOVA followed by Tukey’s multiple comparisons test (to compare multiple treatment groups versus control). For non-normal distribution, the Mann–Whitney U test (two-tailed) was used to compare ALS and the HC cohort. For non-normal multiple comparisons, a Kruskal–Wallis one-way ANOVA (non-parametric) followed by Dunn’s multiple comparisons test was performed (to compare multiple treatment groups versus controls). The diagnostic performance of candidate miRNAs was visualized by receiver operating characteristic (ROC) curves. The area under the curve (AUC) was used to assess the sensitivity and specificity of the candidate biomarkers. Correlations were analyzed by Spearman rank correlation test. All statistical analyses were performed using Prism 9 (GraphPad Software, San Diego, CA, USA). \**P* < 0.05 was considered statistically significant.

### Study approval

This study was conducted under the World Medical Association’s Declaration of Helsinki and approved by the Ethics Committee of Hanyang University Hospital (HYUH IRB 2013-06-012, 2017-01-043). All patients provided written informed consent before inclusion in the study.

## Glossary

ALS(S): slowly progressive ALS; ALS(I): intermediately progressive ALS; ALS(R): rapidly progressive ALS; ALSFRS-R: Amyotrophic Lateral Sclerosis Functional Rating Scale–Revised; AUC: area under the curve; CI: confidence interval; DEGs: differentially expressed genes; ELISA: enzyme-linked immunosorbent assay; HC: healthy control; iMG: microglia-like cell induced by monocyte; NCKAP1: NCK-associated protein 1; NCKAP1L: NCK-associated protein 1 like; NF: neurofilament; NfL; neurofilament light chain; miRNA: microRNA; ROC: receiver operating characteristic; WASP: Wiskott-Aldrich syndrome protein; WAVE: WASP-family verprolin homologous protein; WRC: WAVE regulatory complex

## Author contributions

MY. N and MS. K. designed and performed the experiments, analyzed the data, and wrote the manuscript. KW. O. and JS. P. contributed to the clinical data interpretation. CS. K. and YE. K. performed the genetic analysis. M. N. analyzed the RNA transcripts. JS. B. and HK. J. contributed to the design of the experiments. SH. K. directed the overall study, including designing the experiments, analyzing the data, and writing the manuscript. All authors read and approved the final manuscript.

## Acknowledgments

We thank all staff from the Department of Neurology, College of Medicine, Hanyang University, and the patients for participating in this study. We also thank all members of the Kim Lab for their helpful discussions.

## Ethics approval and consent to participate

This study was conducted under the World Medical Association’s Declaration of Helsinki and approved by the Ethics Committee of Hanyang University Hospital (HYUH IRB 2013-06-012, 2017-01-043).

## Availability of data and materials

All data generated or analysed during this study are included in the source data file and supplementary data.

## Supplementary Materials

Supplemental Table 1. Demographics of the enrolled participants (discovery cohort).

Supplemental Table 2. Demographics of the enrolled participants (validation cohort).

Supplemental Table 3. List of gene panels related to ALS, FTD, and other types of dementia.

Supplemental Table 4. Antibodies, primers, and miRNAs used.

Supplemental Table 5. Plasma cytokine levels in the discovery cohort.

Supplemental Table 6. List of 2,559 shared genes in ALS(R)-iMGs compared to ALS(S)-iMGs according to whole-transcriptome differential gene expression (1.5-fold-change).

Supplemental Table 7. GO analysis of 2,559 genes significantly altered in ALS(R)-iMGs compared to ALS(S)-iMGs.

